# Male age is associated with extra-pair paternity, but not with extra-pair mating behaviour

**DOI:** 10.1101/219477

**Authors:** Antje Girndt, Charlotte Wen Ting Chng, Terry Burke, Julia Schroeder

**Affiliations:** Evolutionary Biology, Max Planck Institute for Ornithology, Seewiesen, Germany; Department of Life Sciences, Imperial College London, Silwood Park Campus, Ascot, United Kingdom; International Max-Planck Research School (IMPRS) for Organismal Biology, University of Konstanz, Konstanz, Germany; Department of Animal and Plant Sciences, University of Sheffield, Sheffield, United Kingdom

**Author notes:** Correspondence: Antje Girndt Julia Schroeder.

**Keywords:** mating behaviour, male manipulation hypothesis, extra-pair paternity, female choice, male age, passerines

## Abstract

Extra-pair paternity is the result of copulation between a female and a male other than her social partner. In socially monogamous birds, old males are most likely to sire extra-pair offspring. The male manipulation and female choice hypotheses predict that age-specific male mating behaviour could explain this old-over-young male advantage. These hypotheses have been difficult to test because copulations and the individuals involved are hard to observe. Here, we studied the mating behaviour and pairing contexts of captive house sparrows, *Passer domesticus*. Our set-up mimicked the complex social environment experienced by wild house sparrows. We found that middle-aged males, that would be considered old in natural populations, gained most extra-pair paternity. However, both female solicitation behaviour and subsequent extra-pair matings were unrelated to male age. Further, copulations were more likely when solicited by females than those initiated by males (i.e. unsolicited copulations), and unsolicited within-pair copulations were more common than unsolicited extrapair copulations. To conclude, our results did not support either hypotheses regarding age-specific male mating behaviour. Instead, female choice, independent of male age, governed copulation success, especially in an extra-pair context and post-copulatory mechanisms might determine why older males sire more extra-pair offspring.

## Introduction

One of the most robust findings in studies of avian extra-pair paternity is that older males sire more extra-pair offspring than younger males (see meta-analyses in^1,2^). What gives older males the competitive edge over younger males is unclear^2^, but the finding has been considered to provide evidence for the ‘good genes’ hypothesis because older males have proven their viability^3^, and are considered to be of high genetic quality (reviewed by^1,4^). Females might seek copulations from older males to obtain genetic benefits for their offspring^5-7^, but see^8,9^. However, there is opposing, albeit inconclusive, empirical evidence for the idea that females gain genetic benefits through extra-pair mating^10-12^.

Extra-pair behaviour involves at least three individuals: the social male, the social female and one extra-pair male^13^. The proximate mechanisms responsible for the positive association of male age with extra-pair paternity are unclear. It has been suggested that older males might outcompete younger males for extra-pair mating opportunities^3-15^ or that females may simply prefer older males as extra-pair partners^16,17^. Alternatively, older males might outcompete younger males post-copulatory through better sperm competition^18^. Here, we test whether older males are better at achieving extra-pair copulations and paternity, and how female solicitation is associated with extra-pair mating.

Weatherhead and Boag (1995) and Westneat and Stewart (2003) suggested that older males are more experienced than younger males and better at convincing or forcing females to mate with them. Hence, older males are predicted to obtain more extra-pair copulations than younger males. This was coined ‘the male manipulation hypothesis’^19^. Through coercive mating, older males are also predicted to achieve more within-pair copulations^13^. Measuring the frequency of extra-pair copulations in wild populations, especially in non-colonial breeding birds, is difficult because extra-pair copulations can be secretive^20^. Several studies have analysed the copulation frequency or display rates of males in relation to their age in birds, e.g.^18,21^ and primates^22,23^. However, we are aware of only one study on the relationship between extra-pair copulations and male age; this showed that extra-pair mating attempts did not correlate with the estimated age of male razorbills, *Alca torda*, (*N* = 15 males)^24^.

The pattern of older males gaining more extra-pair paternity could also be caused by the mating behaviour of the female. The female choice hypothesis is supported by theoretical analysis^25^ but less so by empirical evidence: while a meta-analysis found some support for female birds preernng to copulate with older males^26^, a follow-up review reported mixed results^27^. The female-choice hypothesis is commonly tested by using extra-pair offspring as a proxy, e.g.^2,28^, instead of measuring female choice directly, but see^29^ for a behavioural approach in the wild. This is a limitation because the number of extra-pair offspring reflects only the extra-pair copulations that led to fertilisation, but not how female choice for older extra-pair males is expressed in females behaviourally. For instance, females could either resist extra-pair mating attempts by older males until the costs of resistance are too great, and hence adopt a convenience polyandry strategy *sensu*^13^, or they might actively solicit extra-pair copulations from older males.

We used a captive population of house sparrows, *Passer domesticus*, of known ages to distinguish between those different strategies. We studied the copulation behaviour of both males and females in a semi-natural set-up. House sparrows are particularly suitable to test the predictions of the male manipulation^13,14^ and female choice hypotheses^30^ because, like most passerines, house sparrows are socially monogamous but sexually promiscuous. This means that a male and a female stay together for one, more often multiple, breeding attempt(s)^31^, but copulations with an individual other than the social mate are evident from paternity analyses^11^. Further, male age is the most robust predictor of extra-pair paternity in house sparrows^1,2^. In our set-up, males and females were kept in communal groups to mimic the gregarious colony structures found in wild house sparrow populations^31^. This laboratory environment has the advantage that females can choose among multiple males for within- and extra-pair mating and copulation behaviour can be measured. We first studied (1) the association between extra-pair paternity and male age. We then tested the following predictions from the (2) male manipulation, and (3) female choice hypotheses, and also (4) whether realised extra-pair paternity is a good proxy for copulation behaviour:

(1) We predicted that extra-pair paternity should be positively associated with male age. (2) If older males are better at creating extra-pair opportunities, then we further predict that older males will have more extra-pair copulations. (3) We predict that females solicit more often to older than younger males for both within-and extra-pair copulations and that female solicitation should increase the probability of both within-and extra-pair copulations. Finally, (4) we tested the prediction that the number of extra-pair offspring correlates with extra-pair copulation behaviour.

## Results

### Male age and its association with extra-pair paternity

Across the 400 embryos, 40 were extra-pair (i.e. 10% of all offspring). This value is lower than a recent report on a wild house sparrow population, where 17.5% of all young were extra-pair ^11^. Across broods (*N* = 119), 25 broods contained at least one extra-pair offspring (i.e. 21% of all broods).

We found that extra-pair paternity and male age showed a statistically significant and non-linear relationship in our population: middle-aged males (i.e. 5 years old) sired the highest proportion of extra-pair offspring (Table 1, Fig. 1), e.g. 15% of middle-aged males’ offspring were extra-pair.

**Figure 1:**
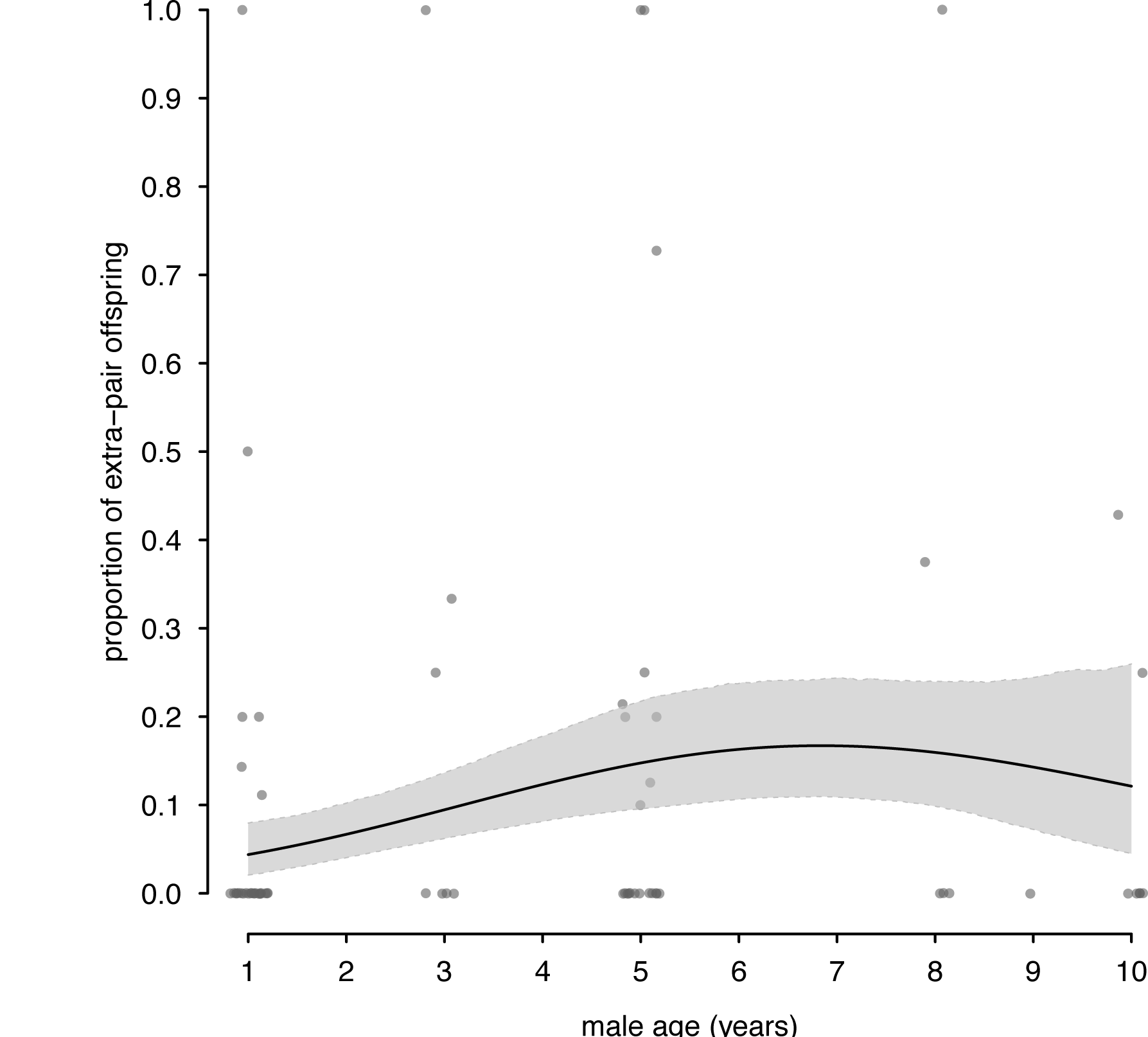
Proportion of extra-pair offspring in relation to the age of male house sparrows, *Passer domesticus* (*N* = 75 males). Middle-aged males sired most extra-pair offspring. We show the average population regression line from the GLM (black line) with CrI (grey area). Open circles represent individual data offset at the x-axis to aid visualization.

**Table 1.**

|  | estimate (lower CrI to upper CrI) |
| --- | --- |
| **Fixed effects** |  |
| (intercept) | -1.50 (-2.11 to -0.88) |
| male age | **0.81 (0.31 to 1.32)** |
| male age<sup>2</sup> | **-0.61 (-1.06 to -0.16)** |
| aviary B | 0.18 (-0.70 to 1.06) |
| aviary C | -0.38 (-1.39 to 0.61) |
| aviary D | **-1.15 (-2.20 to -0.10)** |

The proportion of extra-pair paternity showed a statistically significant quadratic relationship with male age (*N* = 75 males). Results are from a generalised linear model, GLM, assuming a binomial error distribution (logit-link function). Male age was centred and scaled. Extra-pair to within-pair offspring was fitted as a proportional response variable. We show the model’s posterior means and 95% Credible Intervals (CrI). CrIs interpreted as statistically significant are in bold.

### Male manipulation hypothesis

We observed a total of 463 mating attempts, ranging from 0 to 28 per male, and could confirm occurrence of copulation, solicitation as well as the identities of the male and female in 425 of these 463 mating attempts (i.e. 8.3% compromised observations). 107 male mating attempts (23.4%) were directed towards an extra-pair female. Male age did not predict the proportion of extra-pair mating attempts (estimated effect size 0.07 (CrI: −0.19 to 0.33, *N* = 73 males, Fig. 2a, full model output in supplementary information Table S1a). Further, we observed a total of 170 copulations, ranging from 0 to 13 per male. Of these, 27 copulations (19.3%) were with an extra-pair female. Similar to mating attempts, male age did not affect the proportion of extra-pair copulations (estimated effect size 0.03 (CrI: −0.51 to 0.57, *N* = 74 males, Fig. 2b, full model output in supplementary information Table S1b). Additionally, male age was not associated with the total number of mating attempts or copulations (supplementary information Table S2). Notably, 29 of 174 individuals (16.7%, nine males and 20 females) were never observed to be sexually active (i.e. attempting to mate or copulate). Three of these nine sexually inactive males and nine of the 20 sexually inactive females achieved genetic parentage, which means that they copulated unnoticed and represent the subset of individuals that we did not observe.

**Figure 2:**
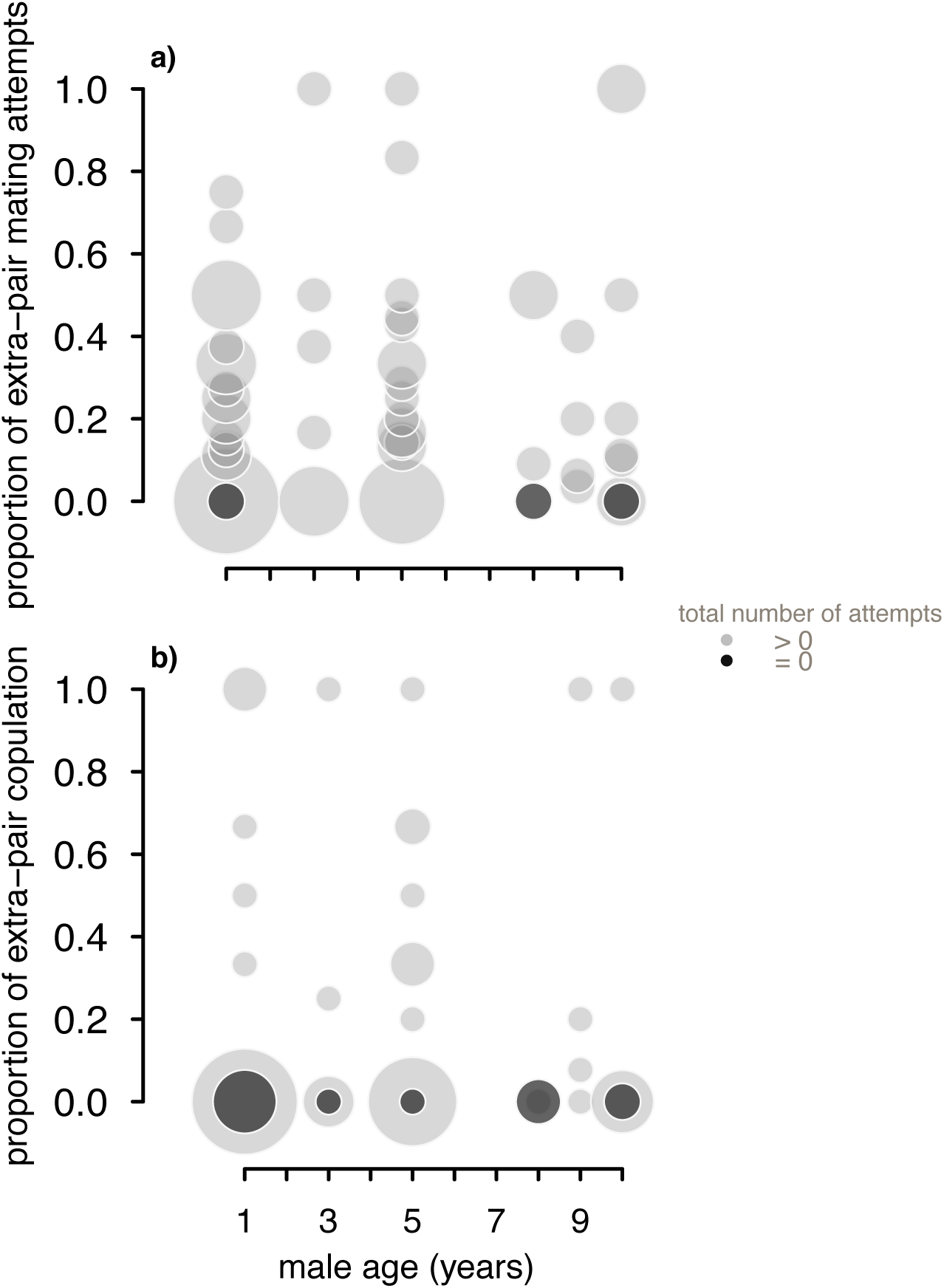
Extra-pair mating behaviour in relation to age in male house sparrows, *Passer domesticus*. Neither the proportion of extra-pair mating attempts (a) (*N* = 73 males) nor the proportion of extra-pair copulations (b) (*N* = 74 males) was explained by the age of males. Circles represent individual data and are scaled according to the number of males of a certain age that were (light grey) or were not observed (dark grey) as sexually active.

### Female choice hypothesis

A within-pair mating attempt was about four-fold and an extra-pair mating attempt about 17-fold more likely to lead to copulation when they were female-solicited compared to mating attempts that were unsolicited (Table 2, Fig. 3). Further, solicited within-pair and extra-pair copulations were equally common but only 4.3% of unsolicited extra-pair mating attempts led to copulation, compared with 19.1% in within-pair matings (Table 2, Fig. 3). The ages of males were not associated with the success of extra-pair or within-pair mating attempts (Table 2). Additionally, the number of unsolicited extra-pair mating attempts was almost double that of solicited extra-pair mating attempts (54 male attempts versus 29 female attempts, binomial test *P* < 0.01, Fig. 3), while the numbers for within-pair mating attempts were more balanced between the sexes (159 male attempts versus 139 female attempts, binomial test *P* = 0.27, Fig. 3).

**Figure 3:**
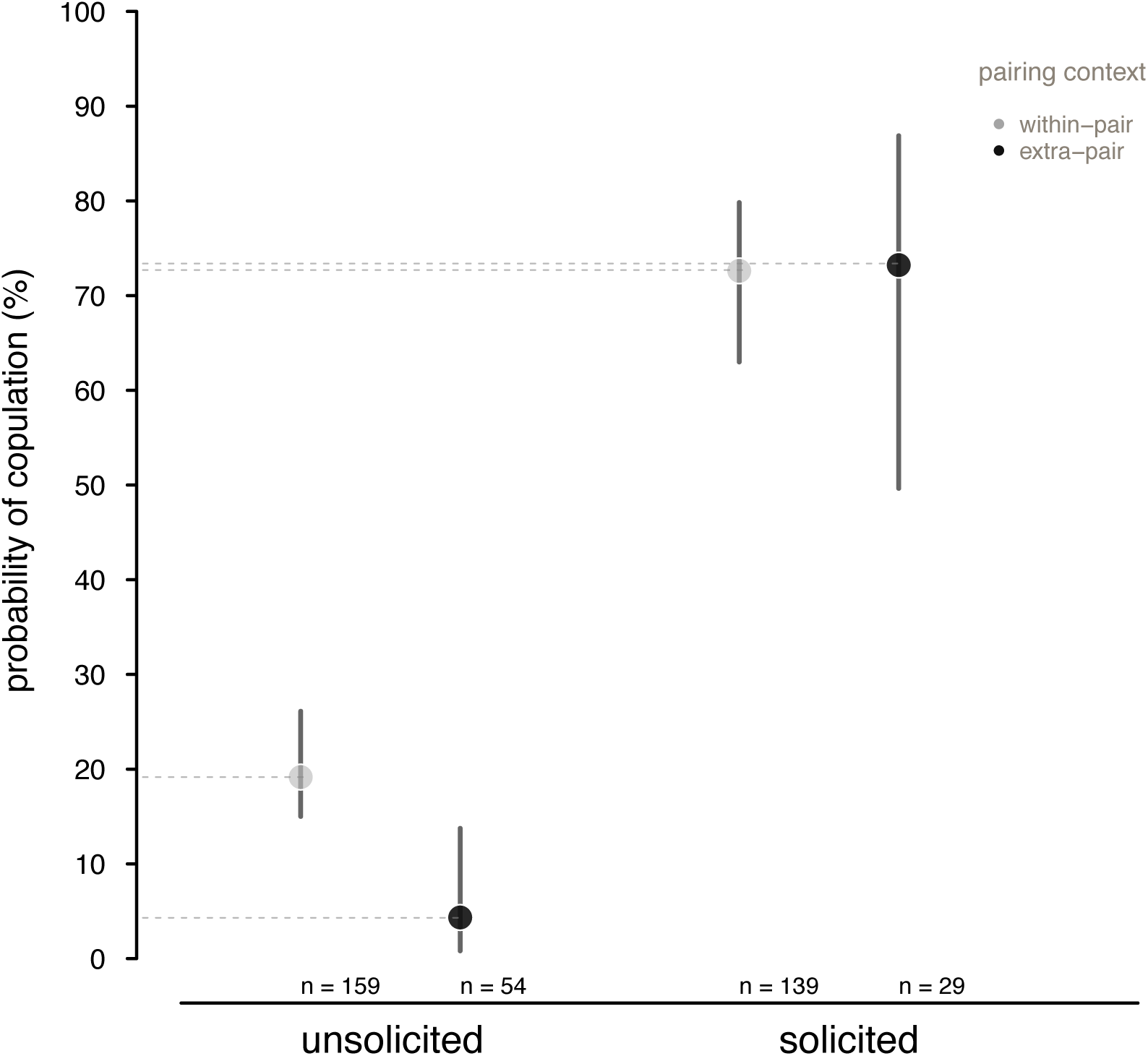
Mating attempts leading to copulation in house sparrows in relation to female solicitation and pairing status (*N* = 381 mating attempts). Female solicitation statistically significantly increased the likelihood of copulation. The effect depended on the pairing context: without female solicitation, copulations were more common with the social male than with an extra-pair male. Unsolicited (i.e. male-initiated) mating attempts were least successful. Filled dots represent posterior model means and the horizontal dashed lines were added to help visualisation. Vertical lines represent CrI.

**Table 2.**

|  | estimate (lower CrI to upper CrI) |
| --- | --- |
| **Fixed effects** |  |
| (intercept) | -1.32 (-1.98 to -0.62) |
| solicited | **2.45 (1.88 to 3)** |
| extra-pair | **-1.87 (-3.45 to -0.28)** |
| male age | -0.03 (-0.36 to 0.30) |
| male age<sup>2</sup> | -0.05 (-0.41 to 0.29) |
| solicited * extra-pair | **1.84 (0.01 to 3.65)** |
| aviary B | -0.27 (-1.04 to 0.54) |
| aviary C | 0.16 ( -0.98 to 0.57) |
| aviary D | -0.19 (-0.98 to 0.56) |
| **Random effects** |  |
| male ID | 0.17 (0.12 to 0.23) |
| female ID | 0 (0 to 0) |

Female solicitation had a statistically significant positive effect on whether a copulation occurred (*N* = 381 mating attempts). In the absence of female solicitation, extra-pair copulations were statistically significantly less common than within-pair copulations. Results are from a GLMM with a binomial error distribution (logit-link function). Female solicitation (‘solicited’, ‘not solicited’) and pairing status (‘within’- or ‘extra-pair’) were categorical fixed effects as well as the interaction of female solicitation and pairing status. Male age was centred and scaled and the outcome variable was a binary response of a mating attempt leading to copulation (‘yes’, ‘no’). We show the model’s posterior means and CrI. CrIs interpreted as statistically significant are in bold.

### Extra-pair offspring as a proxy for extra-pair copulations

The number of extra-pair copulations was not correlated with the number of extra-pair offspring (*N* = 85 males, Spearman rank correlation, *rho* = 0.15, *P* = 0.16, Fig. 4). Of the 85 males in this analysis, 55 males attempted extra-pair mating and 20 subsequently copulated with an extra-pair female compared to 53 out of 85 males that achieved within-pair copulations (see supplementary information Fig. S1 for the correlation of within-pair copulations with within-pair offspring, Spearman rank correlation, *rho* = 0.33, *P* < 0.01). It does not seem reasonable to assume that extrapair correlations correlate as strongly with paternity as within-pair copulations^32^. Still, we cannot exclude the possibility that the lack of correlation reflects missing observations. There was no difference between the average age of males that were observed performing an extra-pair copulation (mean age 4.5 years, *N* = 20 males) and those that were not (mean age 4.6 years, *N* = 65 males, unpaired *t*-test *t*_36.54_ = 0.17, *P* = 0.87).

**Figure 4:**
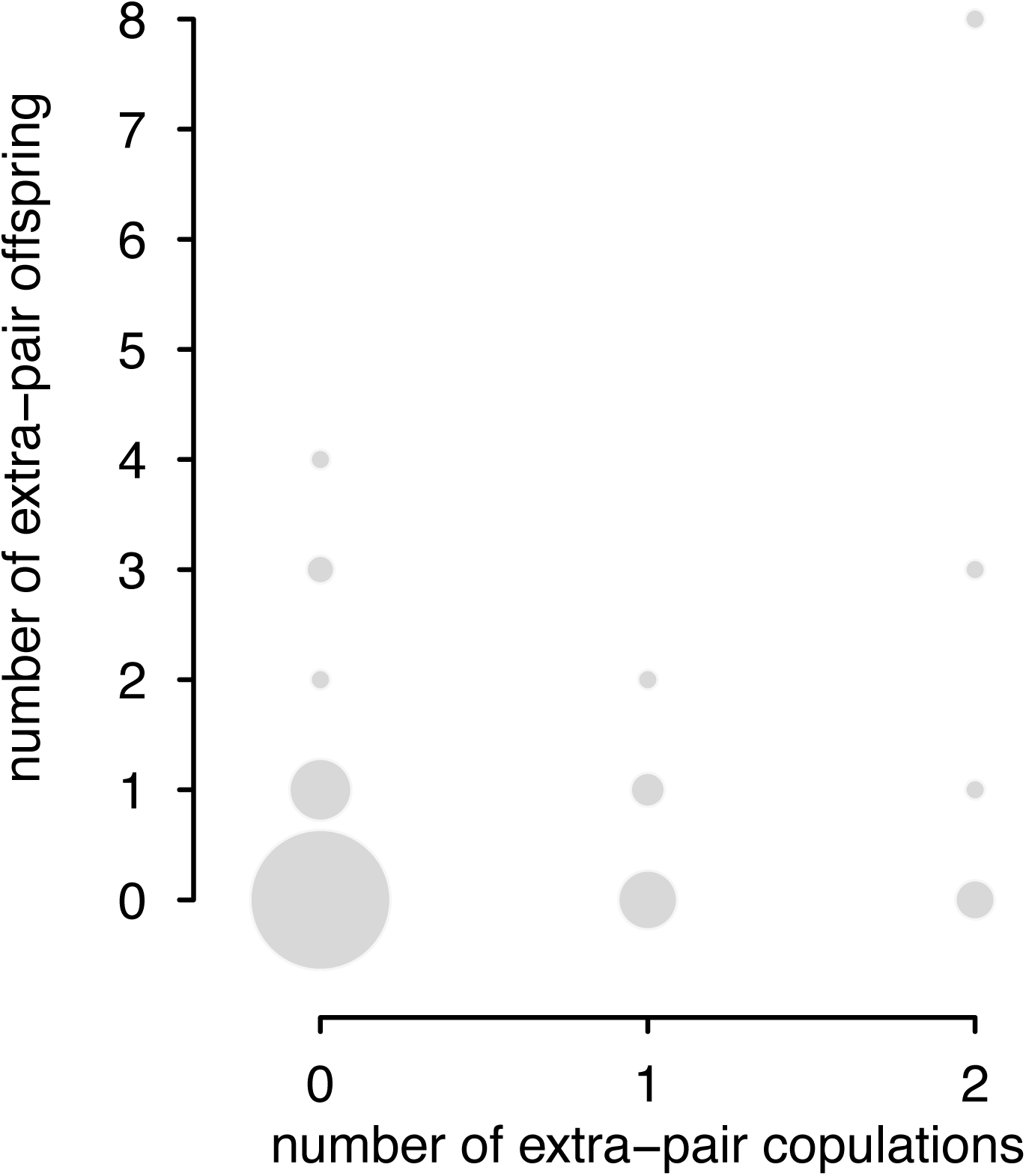
Individual data of extra-pair copulations and extra-pair offspring (*N* = 85 males). The number of extra-pair copulations was not correlated with the number of extra-pair offspring.

## Discussion

In the wild, house sparrows live on average for 3.4 years, and up to a maximum of 13 years^31,33^. Our finding that middle-aged males, old birds in the wild^19,31^, produced most extra-pair offspring mirrors the results in a wild house sparrow population, where extra-pair paternity increased with age in males before showing a decline^19^. The precise age of individuals is known in both studies, which allows extremely old males to be identified and a quadratic age effect on extra-pair paternity to be detected. Further, we did not find an association between extra-pair mating and male age or female choice and male age. Our results imply that male age may not be an important predictor of extra-pair mating behaviour, and our results thus do not support the male manipulation hypothesis 13,14 nor the female choice hypothesis^16,17^.

Male age is the best predictor of extra-pair paternity in wild birds^1^, and our work confirms this in captivity too. Male age and extra-pair mating behaviour, however, were not associated and thus other mechanisms than mating behaviour could drive the relationship between extra-pair paternity and male age. Older males may outcompete younger males via post-copulatory mechanisms; for instance, if older males were better sperm competitors because of larger testes^34^. Alternatively, across iteroparous taxa, individuals show a peak in offspring production before reproductive senescence commences, due to better access to resources^35^ or simply because older individuals have more opportunities to encounter females^36^. As our study used a one-point-in-time sampling approach for all individuals, ensured an equal opportunity for males to encounter females and *ad libitum* access to crucial resources such as nest sites, nesting material and food, the statistically significant non-linear relationship between extra-pair paternity and male age could be the result of post-copulatory mechanisms that favoured fertilisation by older males.

Our study tested for a correlation of extra-pair mating with male age using, to our best knowledge for the first time, a communal breeding set-up of birds of known ages. In our four populations, older males did not attempt nor achieve more extra-pair copulations than younger males. A possible limitation is that a competitive component to an old male mating advantage would have been reduced per individual with our set-up because we increased the number of old males (i.e. our populations did not represent the typical age pyramid found in wild populations: many young and few old individuals). Yet, we predicted an overall effect of male age on extra-pair mating behaviour and reducing the number of males at old ages experimentally would have reduced the chance of detecting a population effect. What might be the most prominent feature of our captive breeding design is the spatial proximity between territories, i.e. nest boxes. Spatial proximity eliminates costs of forays into neighbouring territories and creates opportunities for intrusion that could have elevated the frequency of extra-pair copulations for all males. With close proximity between territories, male pre-copulatory display will also reach multiple females simultaneously, which might further increase the frequency of extra-pair mating behaviour, and the proportion of extra-pair young^38^, but see^39^. However, even if extra-pair mating behaviour had been elevated by our captive set-up, we have still underestimated the number of extra-pair copulations (Fig. 4) but also within-pair copulations (supplementary information Fig. S1).

Proving that females are making an active mate choice is not straightforward^40^. In captivity, choice chamber tests are often used but these do not necessarily reflect female copulation behaviour (see^41^ for a summary). In the wild, extra-pair offspring are used as a proxy, e.g.^28^, but a bias towards older fathers does not necessarily mean that females prefer to copulate with older males. We combined the best of both approaches by allowing females to choose among multiple males of different ages and studying copulation behaviour directly. We found that female solicitation was not associated with male age (supplementary information Table S3). This contrasts with an experimental study where the social mates of western bluebird, *Sialia mexicana*, females were removed: subsequently, females were more likely to accept copulations from intruding males older than their own, absent mate^29^. Differences are anticipated even within species because females will vary in their impetus to copulate promiscuously^42^. Whilst our study does not reveal which traits, if any, females prefer in males^43^, it suggests that male age does not predict whether females solicit copulations or not.

Mating attempts were statistically significantly most likely to succeed when females solicited males. That female cooperation is important for copulations is not surprising in species without intromittent organs^44^. Also, greater female cooperation towards within-pair than to extra-pair mating has been shown before, e.g.^45^ but see^24,46^. Our study takes these findings a step further by showing that the likeliness of a copulation is most dependent on whether it was solicited by a female, not just her cooperation, especially so for extra-pair copulations. We also observed that males, not females, mostly initiated extra-pair mating attempts, which makes sense as the incentive for a female to cheat is lower than for a male^7^, yet only 4.3% of these unsolicited extra pair mating attempts led to copulation. Despite their markedly reduced success, the probability of an unsolicited extra-pair mating succeeding in copulation was between 0.8-14% (see CrI in Fig. 3), which would be enough for selection to act upon, given that the behaviour is linked to reproductive success. In several species, extra-pair copulations have been found to not result in extra-pair paternity^47-49^ but this was not the case in our study. It would be informative to compare the fertilisation efficiency of solicited versus unsolicited extra-pair copulations in future work.

Females actively solicited promiscuous copulations but, in contrast, convenience polyandry, i.e. females giving in to extra-pair males^13^, seems to play a minor role in house sparrows. Female extra-pair behaviour could be explained by indirect selection for alleles that increase male promiscuity^43^. Intriguingly, whilst copulations initiated by females were more successful than those initiated by males, the former were not always successful: approximately 25% of solicitations did not result in a cloacal kiss. There were multiple reasons for mating attempts failing, such as the clumsiness of the couple, or disturbance by conspecifics, which corroborates observations on mating behaviour in zebra finches^43^ but we also witnessed males ignoring female solicitation. A male’s refusal to copulate might be explained by a physiological constraint in house sparrows because males can become ejaculate-depleted within a day^50^. It would be interesting to quantify and better understand the occurrence of male resistance to female mating attempts in future studies.

Our study showed that extra-pair paternity is unlikely to predict extra-pair copulations well, given that male initiated extra-pair mating attempts were mostly unsuccessful. Also, one could expect the relationship between extra-pair copulations and extra-pair paternity to be weaker compared to that between within-pair copulations and within-pair paternity^32^. A lack of a relationship between extra-pair copulations and paternity is, however, biologically implausible and future work should reveal the strength of relationship between extra-pair mating and extra-pair paternity.

To conclude, the observation that females responded more cooperatively to copulation attempts by their social male than by an extra-pair male also emphasises a function of the pair-bond that precedes biparental care. Females also solicited extra-pair copulations, highlighting that extra-pair courtship, despite being male-driven, is a mating strategy adopted by both sexes, the success of which is mainly under female control in house sparrows. Extra-pair copulation will allow some males to increase their reproductive success, and post-copulatory mechanisms might be responsible for the robust correlation between extra-pair paternity and male age in birds.

## Methods

### Study population and experimental breeding set-up

Birds were kept at the Max Planck Institute for Ornithology in Seewiesen, Germany and looked after as described in^51^. The population consisted of wild-caught house sparrows born in 2005 and 2006 and their offspring born in captivity^52^. Males and females were assigned to four aviaries each measuring 3.6 m x 4.0 m x 2.2 m. Per aviary, we had a similar number of males and females, between 21 and 24 pairs, at equal sex ratios and uniform age distributions and there was no evidence for age-assortative mating in our four populations (*N* = 75 social pairs, Spearman rank correlation, *rho* = −0.05, *P* = 0.66). Birds were between one and ten years old but we lacked males aged two, four and six to seven years (Table S4 in the supplementary information gives detail of the age structures and densities per aviary).

House sparrows are hole-nesting passerines that use nest boxes for breeding^53^, so all aviaries were equipped with sufficient individually marked nest boxes to accommodate the respective numbers of pairs plus one extra nest box to reduce competition for sites, e.g. 22 nest boxes for an aviary that held 21 pairs of birds. All birds had *ad libitum* access to food and water^51^, and to nesting material such as hay, horse hair and coconut fibre. Further, each bird was equipped with a combination of a uniquely-numbered metal ring and three coloured plastic rings to allow identification.

### Paternity analysis

Nest boxes were monitored daily. Five to seven days after females initiated incubation, we collected all eggs for parentage analysis, and replaced eggs with fake plaster eggs, resembling house sparrow eggs, to retain natural breeding sequences. We used 12 microsatellite markers^54^ (*Ase18, Pdo1, Pdo3, Pdo5, Pdo6, Pdo9, Pdo10, Pdo16, Pdo17, Pdo19, Pdo22, Pdo27*) and the procedures described in^54^ for genotyping. Cervus version 3.0.7^55^. was used to establish genetic parentage. We first assigned putative mothers from behavioural observations and then, in a second step, we used the confirmed maternity and allowed for all males per aviary to be sires to determine paternity. Of 405 embryos, 400 could be assigned to genetic sires with 95% confidence. For the remaining five embryos, genetic paternity could not be established.

### Behavioural observations

Behavioural observations were made daily from 15 April - 18 June 2015, which represents the beginning and the middle of house sparrow s breeding season^31^. Daily behavioural observations were started between 07:00-07:30 and were recorded live using a Zeiss Victory, 10 x 42 mm, binocular, mostly by CWCT. Three co-workers substituted CWCT on six successive days. Observers were blind with regard to the age and pairing status of individuals when recording birds’ behaviour. As the four aviaries were too large to be observed with an unobstructed view, we divided each aviary into three same-sized sections (see supplementary information Fig. S2 and Fig. S3). Each aviary section was observed separately for 10 to 15 minutes resulting in a total observation time of two to three hours per day. The order of the observations of each aviary section was randomised, using the built-in function sample() in R version 3.2.1^56^, to ensure that observations were not compromised by potential order effects. We identified pair-bonds and nest box owners by later analysing which birds were seen repeatedly at or in each nest box, attending and building nests, and which birds laid and incubated eggs. These criteria were sensible because house sparrows do not engage in pair-bond formation behaviour such as allopreening^31^. Instead, house sparrows commonly initiate pair-bonds after a male has procured a nest site, and the repeated presence of a male and a female at the nest is a strong indication of their pair-bond^31,57^.

We also observed individual copulation behaviour. A male house sparrow displays by approaching a female, lifting his wings slightly, hopping around her vigorously, and vocalising continuously before attempting to mount her^58^. Male house sparrows can also attempt copulation during communal chases of a single female but these chases, while vigorous, rarely result in successful copulations^59^. When females initiate copulation, they adopt a crouching position with their wings quivering and their posterior end held upright (see the video file in the supplementary information). This female behaviour is referred to as solicitation and is distinct from a female’s passive cooperation in a male initiated copulation (i.e. raising her tail and leaning forward to accept a male mating attempt)^31,58^. We refer to a male initiated copulation as an unsolicited copulation in this manuscript. We recorded both a male display and a female solicit, together with the identities of the individuals involved. Subsequently, we recorded whether solicited or unsolicited mating attempts were successful, i.e. resulted in copulation, where a male mounted a female and both individuals bent their tails for a cloacal kiss^60^. In house sparrows, mating behaviour involves copulation bouts comprised of repeated rapid mountings that do or do not include cloacal contact ^31^. The adaptive significance of copulation bouts is not well understood but their occurrence outside the breeding season^59^ highlights that, apart from fertilisation, repeated mounting might be important for pair formation^59^. We used the number of copulation bouts comprising at least one copulation rather than the number of mountings, together with the identities of individuals, in subsequent analyses of whether mating took place within or outside of a pair.

### Ethical note

This study was approved by the Government of Bavaria (Nr 311.5-5682.1/1-2014-024) and the Animal Care and Ethics Committee of the Max Planck Institute for Ornithology.

### Statistical analyses

We used generalised linear models (GLM) and generalised linear mixed effects models (GLMMs,) with a binomial error distribution and a logit-link function to test the questions outlined below. In all models, male age was added as continuous mean centred and scaled explanatory variable, so that the variable male age was measured as the number of standard deviations (sd) from the mean. Aviary identity was included as a fixed effect in all analyses because with only four levels it could not be fitted as a random effect^61^.

#### a) Male age and its association with extra-pair paternity

Using a GLM, we tested whether male age (explanatory variable) positively predicted extra-pair paternity by fitting the number of extra-pair offspring as a proportional response variable (i.e. cbind(number of extra-pair offspring, number of within-pair offspring)). We used a proportional response variable rather than a Poisson GLM because the number of extra-pair offspring was low (overall mean 0.10 extra-pair offspring/offspring) and to adjust for the effect that males that achieve higher paternity inevitably have higher detection rates of extra-pair paternity^62^. As the relationship between extra-pair paternity and male age was expected to be non-linear^19^, we added a quadratic age term as an explanatory variable to the model. We excluded 11 males that were unpaired and thus could be considered floaters^63^. However, qualitatively, the results remained similar to when floaters were included (supplementary information Table S5). The total sample size, excluding floaters, was 75 males.

#### b) Male manipulation hypothesis

Here, we assessed whether male extra-pair mating behaviour was positively associated with male age (explanatory variable fitted as both a linear and quadratic age term) by using two proportional response variables. The first response variable was the proportion of a male's extra- to within-pair mating attempts (i.e. cbind(number of extra-pair mating attempts, number of within-pair mating attempts)). We excluded two outliers that caused overdispersion^64^ but first established that this decision did not mediate our analysis by re-running the analysis including the two outliers and confirming that the results were qualitatively similar. The second response variable was the proportion of a male's extra-pair to within-pair copulations (i.e. cbind(number of extra-pair copulations, number of within-pair copulations)). For both analyses, we again excluded 11 male floaters^63^ but the results remained similar when floaters were included (supplementary information Table S6). Also, four males were paired to two females simultaneously, they were socially polygynous. For the latter males, we summed the mating attempts and copulations for both their pairbonds and only considered mating attempts and copulations outside their two pairbonds as extra-pair. The total sample size, without floaters, was 75 males for the mating attempts GLM and 74 males for the copulation GLM.

#### c) Female choice hypothesis

To assess how female choice affects the likelihood of copulation, we fitted the probability of whether a mating attempt led to copulation (‘yes’ or ‘no’) as a response variable in a GLMM. Female solicitation (‘solicited’, ‘not solicited’) and pairing context (‘within’- or ‘extra-pair’) were categorical explanatory variables as well as the interaction between both. Male age was added as an explanatory variable, including a quadratic age term. Having both female solicitation and male age as predictors in the same model was justified because there was no association between male age and female solicitation behaviour (supplementary information Table S3), which implies that the effects can be interpreted independently from each other and the analyses did not suffer from collinearity. We excluded five floaters present in this dataset, but the analysis including male floaters yielded similar results (supplementary information Table S7). Again, only mating attempts and copulations outside both pair-bonds for socially polygynous males were considered to be extra-pair. The total sample size was 381 copulations attempts involving 71 males, excluding floaters. As repeated measures were obtained across males and females, male and female IDs were added as a random intercept.

#### d) Extra-pair offspring as a proxy for extra-pair copulations

Finally, we tested whether the number of observed extra-pair offspring was correlated with the number of observed extra-pair copulations using a Spearman rank correlation test.

We used R version 3.4.1^56^ and the package “lme4“^65^ to run models. We then used the package ‘’arm’’ and the function ‘’sim’’ to simulate values from the posterior distributions (*n* = 2000 draws) of the model parameters from lme4, assuming non-informative priors. From the simulated values, we extracted 95% Credible Intervals (CrI), based on the 2.5% and 97.5% quantiles of the posterior distributions^66^. The Cr represent the uncertainty of our estimates but we also used them for statistical significance testing because CrI not overlapping zero can be interpreted as a Frequentist p-value < 0.05^64^. For all models, we followed the recommendation in to ensure that model assumptions and fit were met, including checking for overdispersion and multi-collinearity.

## Data Availability

All datasets are available at the Open Science Framework

(http://dx.doi.org/10.17605/OSF.IO/FYURG).

## Acknowledgements

We are grateful to A. Grötsch and N. Fischer for animal care, B. Kempenaers for logistical support and N. dos Remedios for molecular work. We thank A. Sánchez-Tójar for comments on the manuscript. This work was supported by the German Research foundation (DFG grant SCHR 1447/1-1 to JS) and the Natural Environment Research Council (NERC Grant NE/J024597/ to TB).

## Author Contributions

J.S. and A.G. conceived the study. C.W.T.C. and A.G. collected the primary data and T.B. provided for molecular work. A.G. analysed the data, prepared the figures and wrote the paper with input from all the authors.

## Additional Information

The authors declare no competing financial interests.

## SUPPLEMENTARY INFORMATION

**Table S1.**
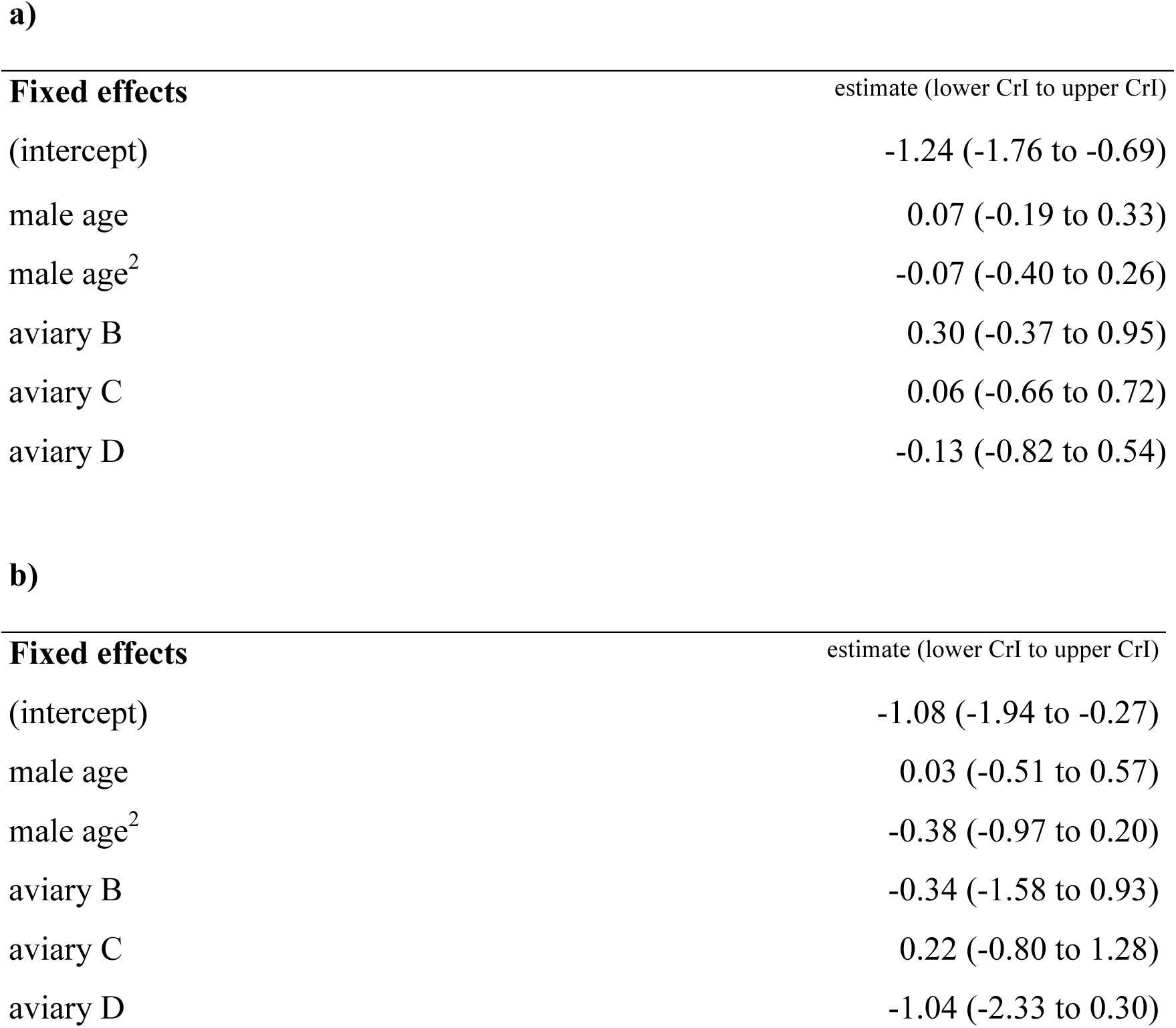

Neither the proportion of extra-pair mating attempts (a) (*N* = 73 males) nor the proportion of extra-pair copulations (b) (*N* = 74 males) was explained by the age of male house sparrows, *Passer domesticus*, excluding floaters. Results are from a GLM, assuming a binomial error distribution (logit-link function). Male age was centred and scaled. A) Extra- to within-pair mating attempts and b) extra- to within-pair copulations were fitted as a proportional response variable. We show the model’s posterior means and CrI. CrIs interpreted as statistically significant are in bold.

**Table S2.**

| **Fixed effects** | estimate (lower CrI to upper CrI) |
| --- | --- |
| (intercept) | 1.76 (1.52 to 1.98) |
| male age | -0.04 (-0.15 to 0.08) |
| male age<sup>2</sup> | -0.07 (-0.20 to 0.06) |
| aviary B | -0.12 (-0.39 to 0.18) |
| aviary C | 0.11 (-0.16 to 0.38) |
| aviary D | 0.17 (-0.10 to 0.43) |

b)
| **Fixed effects** | estimate (lower CrI to upper CrI) |
| --- | --- |
| (intercept) | 0.87 (0.50 to 1.21) |
| male age | -0.02 (-0.19 to 0.17) |
| male age<sup>2</sup> | -0.12 (-0.34 to 0.10) |
| aviary B | **-0.63 (-1.17 to -0.09)** |
| aviary C | 0.13 (-0.31 to 0.56) |
| aviary D | 0.25 (-0.18 to 0.66) |

Neither the total number of mating attempts (a) (*N* = 73 males) nor the total number of copulations (b) (*N* = 74 males) was explained by the age of male house sparrows, excluding floaters. Results are from a GLM, assuming a Poisson error distribution (log-link function). Male age was centred and scaled. We show the model’s posterior means and CrI. CrIs interpreted as statistically significant are in bold.

**Table S3.**

|  | estimate (lower CrI to upper CrI) |
| --- | --- |
| **Fixed effects** |  |
| (intercept) | -0.16 (-0.75 to 0.44) |
| male age | 0.13 (-0.16 to 0.43) |
| male age<sup>2</sup> | -0.13 ( -0.44 to 0.18) |
| aviary B | -0.56 (-1.32 to 0.12) |
| aviary C | 0.09 (-0.64 to 0.84) |
| aviary D | 0.31 (-0.39 to 1.07) |
| **Random effects** |  |
| male ID | 0.42 (0.30 to 0.54) |

Male age (*N* = 77 males) did not explain the probability of solicitation (449 observations) in house sparrows, excluding floaters. Results are from a generalised linear mixed effect model, GLMM, assuming a binomial error distribution (logit-link function). Male age was centred and scaled and the outcome variable was a binary response of solicitation (‘yes’, ‘no’). We show the model’s posterior means and CrI. CrIs interpreted as statistically significant are in bold.

**Table S4.**

|  | aviary A |  | aviary B |  | aviary C |  | aviary D |  |
| --- | --- | --- | --- | --- | --- | --- | --- | --- |
|  | males | females | males | females | males | females | males | females |
| age in years | *N*= | *N*= | *N*= | *N*= | *N*= | *N*= | *N*= | *N*= |
| 1 | 8 | 8 | 8 | 9 | 8 | 5 | 7 | 6 |
| 2 | 0 | 0 | 0 | 0 | 0 | 2 | 0 | 0 |
| 3 | 2 | 2 | 2 | 5 | 2 | 4 | 3 | 5 |
| 5 | 6 | 6 | 6 | 4 | 4 | 4 | 5 | 5 |
| 8-10 | 5 | 5 | 8 | 6 | 7 | 6 | 6 | 5 |
| total number | 21 | 21 | 24 | 24 | 21 | 21 | 21 | 21 |

We aimed at similar sample sizes per age and sex in our four house sparrow populations: young breeders (one to three years old), middle-aged breeders (five-year-old) and old breeders (eight to ten years old).

**Table S5.**

|  | estimate (lower CrI to upper CrI) |
| --- | --- |
| **Fixed effects** |  |
| (intercept) | -1.29 (-1.88 to -0.70) |
| male age | **0.69 (0.24 to 1.15)** |
| male age<sup>2</sup> | **-0.57 (-1.04 to -0.11)** |
| aviary B | -0.01 (-0.90 to 0.90) |
| aviary C | -0.58 (-1.61 to 0.43) |
| aviary D | **-1.33 (-2.36 to -0.30)** |

The proportion of extra-pair paternity in relation to the age of male house sparrows, including unpaired males, i.e. floaters^1^, showed a significant quadratic relationship with male age (*N* = 86 males). Results are from a generalised linear model, GLM, assuming a binomial error distribution (logit-link function). Male age was centred and scaled. Extra-pair to within-pair offspring was fitted as a proportional response variable. We show the model’s posterior means and 95% Credible Intervals (CrI). CrIs interpreted as statistically significant are in bold.

**Table S6.**

|  | estimate (lower CrI to upper CrI) |
| --- | --- |
| **Fixed effects** |  |
| (intercept) | -1.21 (-1.74 to -0.68) |
| male age | 0.16 (-0.09 to 0.41) |
| male age<sup>2</sup> | 0.01 (-0.31 to 0.31) |
| aviary B | 0.23 (-0.44 to 0.91) |
| aviary C | 0.05 (-0.59 to 0.73) |
| aviary D | -0.15 (-0.81 to 0.52) |

**b)**
|  | estimate (lower CrI to upper CrI) |
| --- | --- |
| **Fixed effects** |  |
| (intercept) | -1.08 (-1.93 to -0.24) |
| male age | 0.03 (-0.50 to 0.59) |
| male age<sup>2</sup> | -0.38 (-0.98 to 0.18) |
| aviary B | -0.34 (-1.63 to 0.99) |
| aviary C | 0.22 (-0.90 to 1.28) |
| aviary D | -1.04 (-2.35 to 0.23) |

Neither the proportion of extra-pair mating attempts (a) (*N* = 84 males) nor the proportion of extra-pair copulations (b) (*N* = 85 males) was explained by the age of male house sparrows, including floaters. Results are from a GLM, assuming a binomial error distribution (logit-link function). Male age was centred and scaled. A) Extra- to within-pair mating attempts and b) extra- to within-pair copulations were fitted as a proportional response variable. We show the model’s posterior means and CrI. CrIs interpreted as statistically significant are in bold.

**Table S7.**

| **Fixed effects** | estimate (lower CrI to upper CrI) |
| --- | --- |
| (intercept) | -1.32 (-2 to -0.64) |
| solicited | **2.44 (1.88 to 2.98)** |
| extra-pair | **-1.94 (-3.45 to -0.46)** |
| male age | -0.04 (-0.38 to 0.28) |
| male age<sup>2</sup> | -0.04 (-0.40 to 0.29) |
| solicited * extra-pair | **1.77 (0 to 3.53)** |
| aviary B | -0.27 (-1.08 to 0.60) |
| aviary C | -0.02 (-0.83 to 0.82) |
| aviary D | -0.19 (-0.94 to 0.66) |
| **Random effects** |  |
| male ID | 0.22 (0.16 to 0.30) |
| female ID | 0 (0 to 0) |

Female solicitation had a significant positive effect on whether a copulation occurred in house sparrows, including floaters (*N* = 391 mating attempts). In the absence of female solicitation, extra-pair copulations were significantly less common than within-pair copulations. Results are from a GLMM with a binomial error distribution (logit-link function). Female solicitation (‘solicited’, ‘not solicited’) and pairing status (‘within’- or ‘extra-pair’) were categorical fixed effects as well as the interaction of female solicitation and pairing status. Male age was centred and scaled and the outcome variable was a binary response of a mating attempt leading to copulation (‘yes’, ‘no’). We show the model’s posterior means and CrI. CrIs interpreted as statistically significant are in bold.

**Figure S1.**
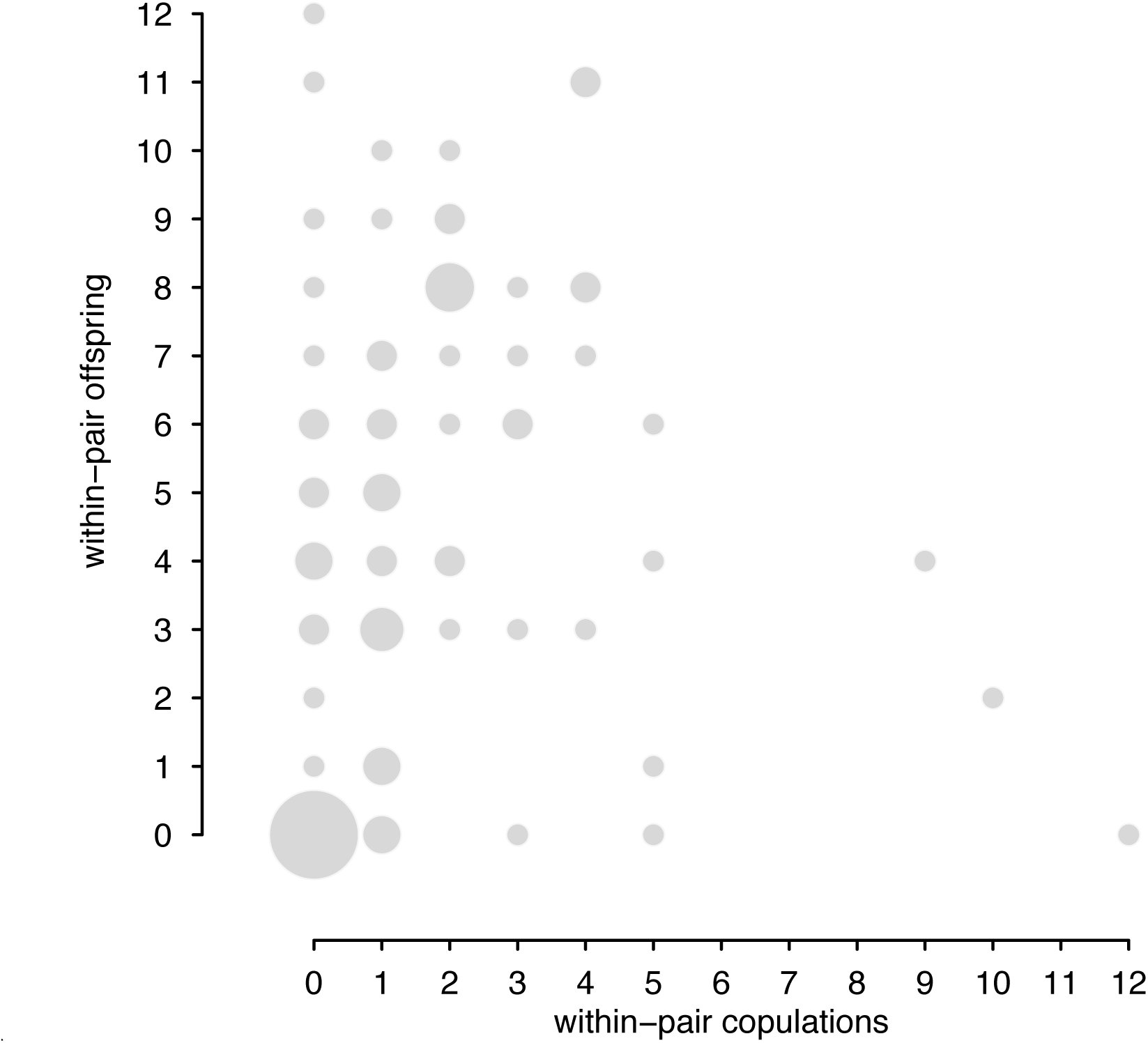
Individual data of within-pair copulations and within-pair offspring (*N* = 85 males). The number of within-pair copulations was correlated with the number of within-pair offspring (Spearman rank correlation, *rho* = 0.33, P < 0.01)

**Figure S2.**
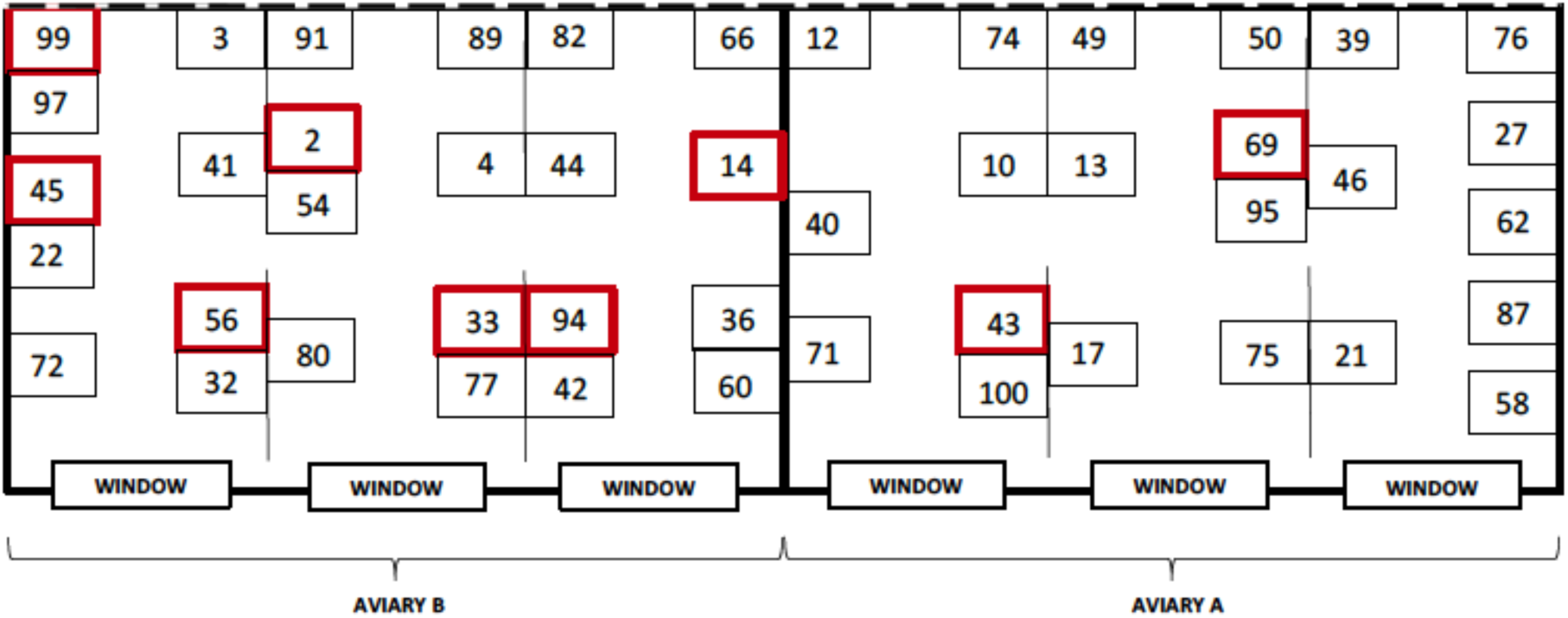
Schematic unscaled bird’s eye view of two of the four house sparrow aviaries. Numbered squares illustrate nest boxes. Red-bordered squares are nest boxes that were fitted above each other, displaced by 30 cm. For example, in aviary B, nest box 99 was fitted 30 cm above nest box 97. Vertical bold lines represent aviary walls. Vertical interrupted lines highlight single open sections within each aviary. The dashed lines represent the outer-wall that was covered with mesh wire. Observations were performed daily through the window into each individual section over a period of three months during the breeding season.

**Figure S3.**
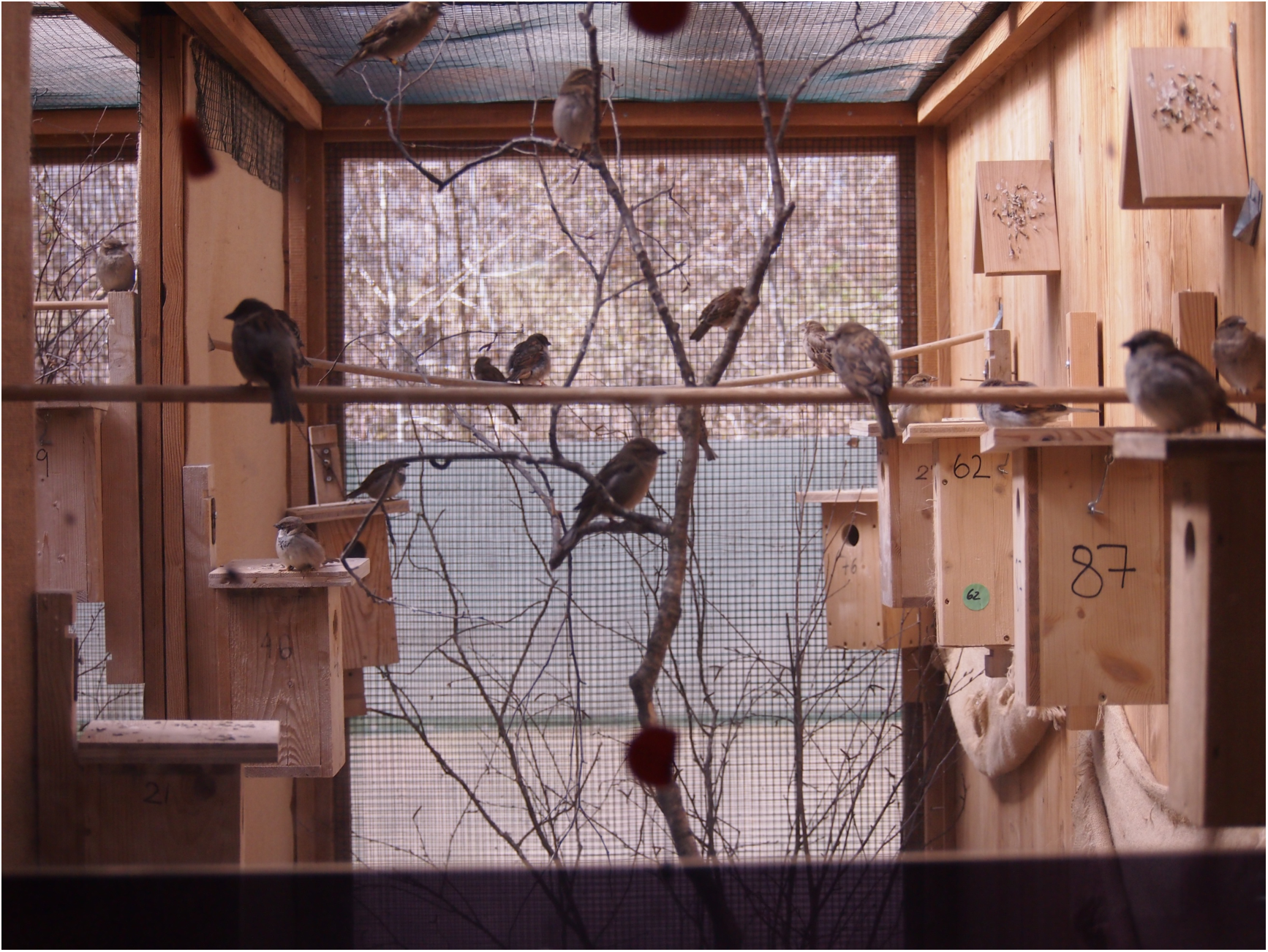
Example view of one house sparrow aviary section. Observations were performed in close proximity to the aviary section window but in contrast to the photograph the observer could see the whole aviary section and not just the upper part.

